# Graph-Theoretical Formulation of the Generalized Epitope-based Vaccine Design Problem

**DOI:** 10.1101/845503

**Authors:** Emilio Dorigatti, Benjamin Schubert

## Abstract

Epitope-based vaccines have revolutionized vaccine research in the last decades. Due to their complex nature, bioinformatics plays a pivotal role in their development. However, existing algorithms address only specific parts of the design process or are unable to provide formal guarantees on the quality of the solution. Here we present a unifying formalism of the general epitope vaccine design problem that tackles all phases of the design process simultaneously and combines all prevalent design principles. We then demonstrate how to formulate the developed formalism as an integer linear program which guarantees optimality of the designs. This makes it possible to explore new regions of the vaccine design space, analyze the trade-offs between the design phases, and balance the many requirements of vaccines.

## Introduction

In recent years vaccines based on T-cell epitopes, so called epitope-based vaccines (EV), have become wildly used as therapeutic treatments in case of cancer immunotherapy [1–4] and prophylactically against infectious diseases [5–10]. Compared to regular attenuated vaccines, EVs offer several advantages [11]. Since EVs are based on small peptide sequences, they can be rapidly produced using well established technologies and easily stored freeze-dried [11]. EVs also do not bare the risk of reversion to virulence as they do not contain any infectious material, and the selection of epitopes can be tailored to address the genetic variability of a pathogen and that of a targeted population or individual increasing its potential efficacy [11].

To aid the design process, bioinformatics approaches have been developed to (1) discover potential candidate epitopes, (2) select a set of epitopes for vaccination, and (3) assemble the selected epitopes into the final vaccine (Figure 1A). Most of the proposed selection and assembly approaches focus either on peptide cocktail vaccines (Figure 1A(3a)) or on so-called string-of-beads vaccines (Figure 1A(3b)), which are polypeptides connecting each epitope directly or by short spacer sequences. Vider Shalit *et al.* for example developed a genetic algorithm that selects epitopes to maximize the coverage of viral and human variation while simultaneously optimizing the ordering of the string-of-beads to increase efficacy [12]. Toussaint *et al.* proposed an approach that selects a fixed number of epitopes to maximize vaccine immunogenicity using integer linear programming (ILP) [13], and later established a method to find the optimal string-of-beads ordering based on a traveling salesperson problem (TSP) embedding [14], which has been recently extended by Schubert *et al.* to incorporate optimal spacer sequences as well [15]. Lundegaard *et al.* proposed a greedy algorithm for epitope selection to maximize antigen and population coverage using a sub-modular function formulation [16].

**Figure 1:**
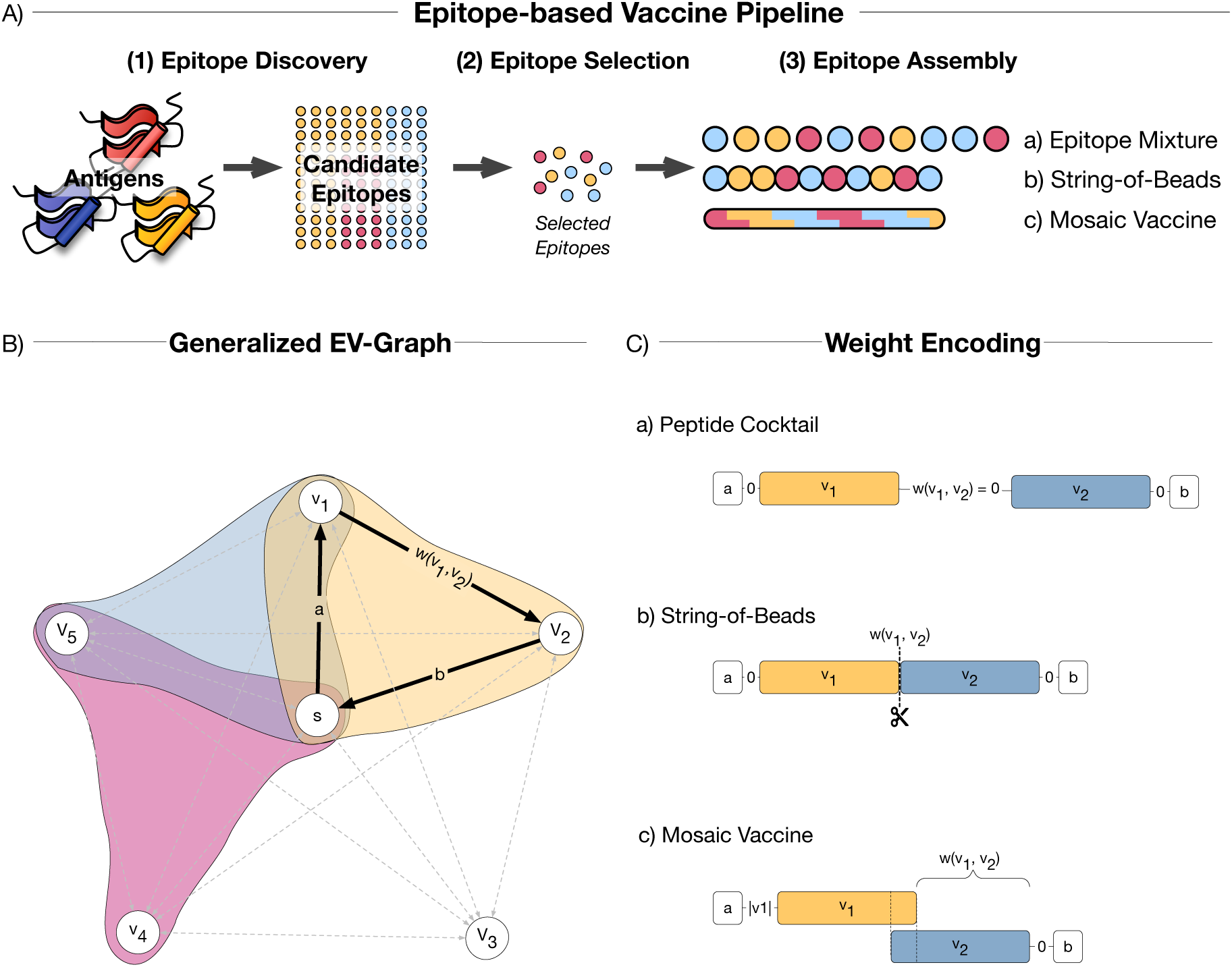
(A) The rational vaccine design pipeline. A epitope-based vaccine pipeline is comprised of three major steps (1) epitope discovery, (2) selection, and (3) assembly. (B) The graph encoding the vaccine design problem. Vertices represent epitopes and edge weights are design specifically defined. By jointly modeling the epitope selection and vaccine assembly problem, we seek a subset of vertices with the highest immunogenicity whose simple tour is not larger than a upper limit. (C) Graphical illustration of the weights used in the three designs. The edge weights are simply ignored for epitope mixtures. For string-of-beads, the edge weight represents the negative log-likelihood of being cleaved at the junction site of the two connecting epitopes. The weights in mosaic designs represents the added length to the mosaic vaccine once joining the two connecting epitopes at their overlap.

Recent studies suggest that through the usage of artificial proteins of overlapping epitopes, so-called mosaic vaccines (Figure 1A(3c)), both depth and breadth of the T-cell response can be remarkably increased [5–10, 17]. Mosaic vaccines constitute an interesting alternative to string-of-beads EVs, as they incorporate many more epitopes within the same vaccine length [18]. This is especially useful for vaccine development against highly polymorphic viruses like Influenza or HIV. A single mosaic vaccine can be designed to cover the observed variability of the virus by targeting multiple antigens, thereby increasing the potential of obstructing virus escape pathways [7]. To aid the design of such mosaic vaccines, Fischer *et al.* introduced a genetic algorithm that constructs a mosaic protein maximizing the number of nine-mer peptides of an antigen pool [19].

Although multiple algorithms exist to aid EV design, they either lack a theoretical foundation, or only model a sub-problem of the entire design problem. Algorithms in the former category cannot provide any guarantees on the quality of the solution and can be arbitrarily far away from the optimal one, which makes comparisons of different designs potentially unreliable. Algorithms in the latter category, in contrast, are unable to capture the trade-offs between different design stages, thus limiting the space of EV design that can be explored.

In this work, we, therefore, develop a rigorous mathematical formulation that models the entire design process, from epitope selection to assembly, unifies all EV design principles, and can be solved to optimality, at the expense of potentially high computational cost. We then use this framework to explore the trade-off between optimal epitope selection and optimal epitope assembly for string-of-beads vaccines, design a cocktail of polypeptides that have the same properties of a much longer vaccine, and show the advantages of mosaic over string-of-beads vaccines in terms of immunogenicity, coverage, and conservation. Finally, we give recommendations on which settings to tune for designing effective mosaic vaccines, and how to reduce the computational burden with little compromise on the quality of the solution.

## Materials and methods

### Graph theoretical formalism combines all EV design principles

The simplest design principle – epitope mixture vaccines – seeks to find a subset *P* of *k* epitopes that together have the highest chance of invoking an effective immune response *I*(*P*). Similarly, the string-of-beads design problem seeks to find a polypeptide comprised of *k* concatenated epitopes that simultaneously maximize the vaccine efficacy *I*(*P*) and the recovery likelihood of each epitope by the proteasome, which is influenced by the ordering of the epitopes in the construct [20]. In contrast, the mosaic design problem is concerned with constructing an artificial antigen *P* of fixed length *h* comprised of potentially overlapping epitopes with maximal efficacy *I*(*P*). These design principles can be further generalized to allow the composition of a cocktail of several poylpeptides that jointly optimize the vaccine efficacy, thereby reducing the overall length of the fragments without sacrificing other properties of the vaccine.

These three design principles can be unified under a single mathematical framework (Figure 1B and C). We formulate the generalized EV design problem as a combinatorial optimization problem on a weighted, directed graph *G*(*V, E, w*), where the vertices *V* represent the epitopes and the the weight *w*(·) of the edges *E* determine the design of the EV. We also add an artificial node *s* representing the N- and C-terminus of the vaccine, connecting it to every vertex *v* ∈ *V* such that *w*(*e*_*sv*_) = *a* and *w*(*e*_*vs*_) = *b*, with design-dependent weights *a, b* ∈ ℝ. To find the optimal EV in 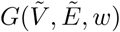, with 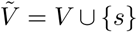 and 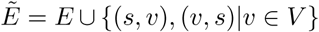, we are seeking *n* disjoint subsets 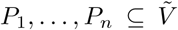, each of size at most *k*, that together maximize the vaccine’s immunogenicity *I* : 2^*V*^ → ℝ, and whose simple tours *H*(*P*_1_), …, *H*(*P*_*n*_) start and end at 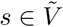 and weigh at most *h* ∈ ℝ (Eq. 1). Here we use the term *simple tour* to refer to a closed walk with no repeated vertices except for *s*.

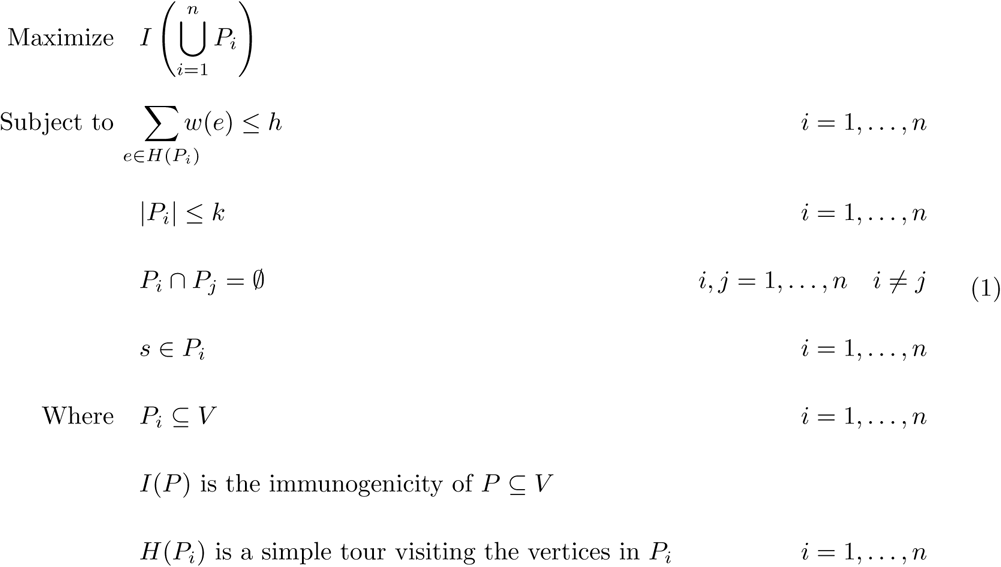

Similar constrained design problems appear in genome assembly [21], and have been extensively studied in the field of operations research under the name of price collecting traveling salesperson, bank robber, or team orienteering problem [22]. Importantly, they are known to be NP-hard [23].

While it is unclear what a good vaccine immunogenicity function *I*(*P*) constitutes, we only will define vaccine efficacy vaguely as the ability to induce a broad immunization in a target population represented by a set of prevalent HLA molecules against a given polymorphic antigen pool, and refer to immunogenicity functions introduced in the context of rational vaccine design [12, 13, 16, 17]. However, for the purpose of algorithmic analysis and comparison, we use the function proposed by Toussaint *et al.* [13], which assumes that each epitope contributes independently to the vaccine’s overall immunogenicity with respect to the target population represented by a set of HLA alleles:

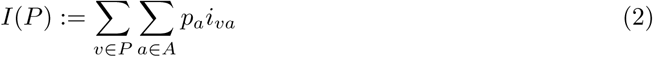

where *p*_*a*_ is the observed frequency of HLA allele *a* ∈ *A* within the target population and *i*_*va*_ is the individual immunogenicity generated by epitope *v* bound to HLA molecule *a*. We approximate the latter with the log-transformed IC_50_ binding strength between the epitope and the MHC complex, which can be predicted by machine learning algorithms such as NetMHCpan [24].

#### Adaptations for epitope mixture design

To design epitope mixture vaccines with the proposed framework, we ignore the edge weight constraint by setting *h* = ∞, and the edge weights *w*(*e*_*ij*_) = 0 for 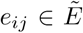 (Figure 1C(a)). The size *k* of *P*, however, has to be defined. Note that this is equivalent to the framework proposed by Toussaint *et al.*[13].

#### Adaptations for string-of-beads design

To enable string-of-beads designs with the framework, we interpret the edge weights *w*(*e*_*ij*_) as the negative proteasomal cleavage log-likelihood between epitope *v*_*i*_ and *v*_*j*_, and set the out- and in-going edges of node *s* to *w*(*e*_*sv*_) = *w*(*e*_*vs*_) = 0 for every *v* ∈ *V* (Figure 1C(b)), following [14]. The proteasomal cleavage likelihood of an epitope can be predicted with existing proteasomal cleavage site methods such as ProteaSMM [25], PCM [26], or NetChop [27].

Solving the so-defined generalized EV design problem yields the string-of-beads design with maximal immunogenicity, whose overall cleavage likelihood is at least *h*. Predetermining *h*, however, is difficult. We might not even be interested in a solution with a fixed *h*, but rather want to explore the inter-dependencies between the immunogenicity objective and the overall cleavage likelihood of the string-of-beads EV. This leads to a reinterpretation of the design formulation as bi-objective optimization problem, in which we simultaneously optimize the overall immunogenicity *I*(*P*) and the length of the tours *H*(*P*_*i*_), *i* = 1, …, *n* (i.e., the overall cleavage likelihood). We can then explore the Pareto frontier of this problem with methods such as the augmented *E*-constraint [28] (section A in Supplement 1).

#### Adaptations for mosaic design

For mosaic vaccines, we define the edge weight *w*(*e*_*ij*_) as the length that would be added to the mosaic antigen once *v*_*i*_ and *v*_*j*_ are joined at their longest suffix-prefix overlap (Figure 1C(c)):

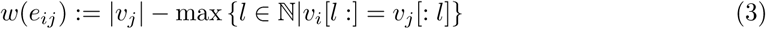

where |*v*_*j*_| represents the length of the epitope sequence *v*_*j*_ and *v*_*i*_[*l* :], *v*_*j*_[: *l*] represent the substrings of length *l* of epitopes *i* and *j* starting and ending at position *l* respectively. Note that Eq. 3 can be computed efficiently in time *O*(*m* + *k*^2^) for a given set of *k* strings of total length *m* by using generalized suffix trees [29]. Furthermore, we define the weights of the out- and in-going edges of node *s* as *w*(*e*_*sv*_) = |*v*| and *w*(*e*_*vs*_) = 0 for all *v* ∈ *V*. The edge-weight sum of any tour that starts and ends at *s* is then equal to the length in amino acids of the resulting mosaic sequence.

### Formulation as an integer linear program guarantees optimally

With the aforementioned definitions, we can formulate the generalized EV design problem as an integer linear program (ILP) encoding the team orienteering problem [22]. This guarantees to construct an optimal EV with the cost of potentially long run times and/or memory requirements, since the number of variables and constraints grows quadratically with the number of epitopes in consideration, usually in the order of 10^3^ or 10^4^.

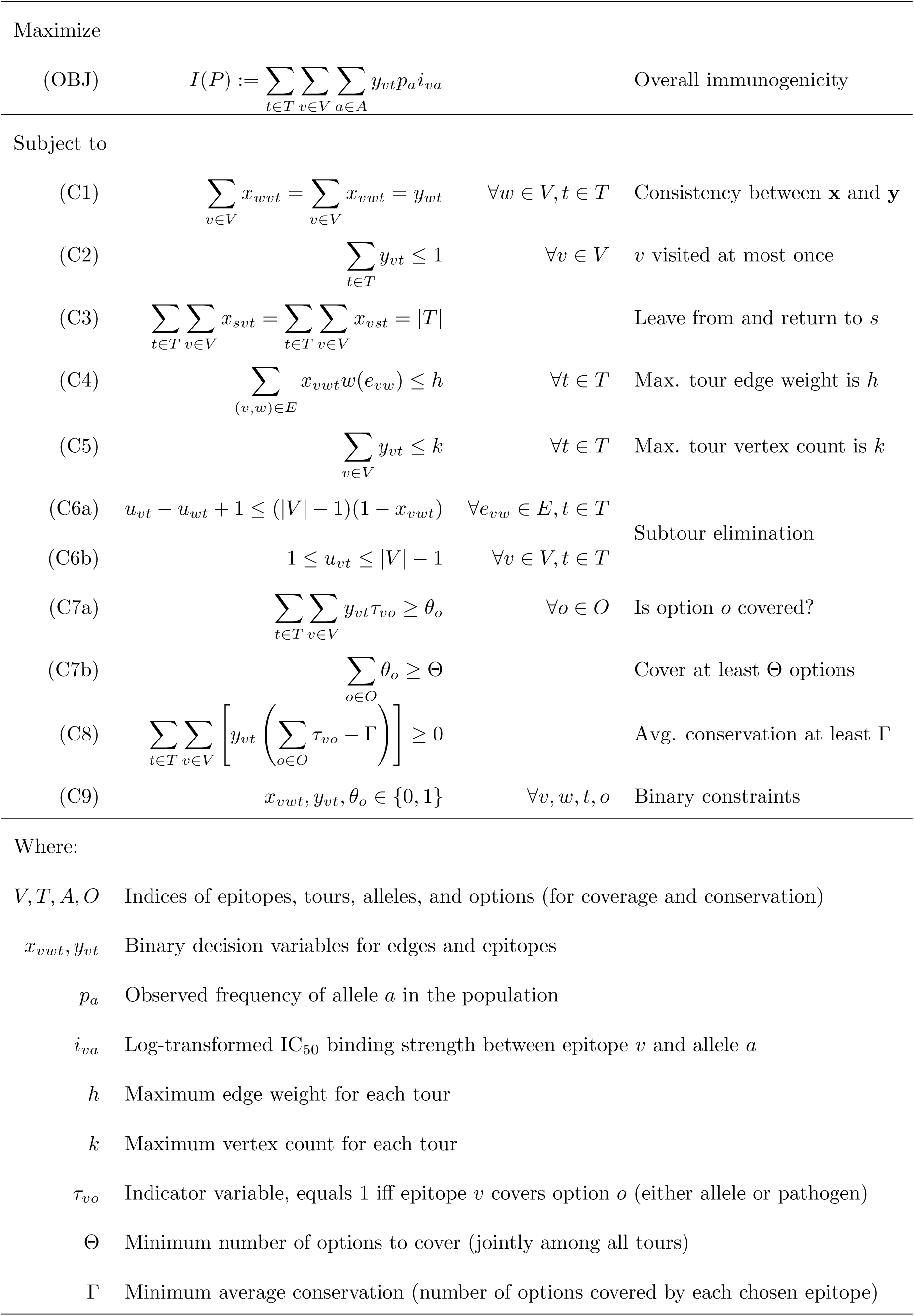

Every tour represents a single polypeptide. We introduce binary decision variables *x*_*vwt*_ and *y*_*vt*_ indicating whether the tour *t* is visiting the edge between *v* and *w* and the vertex *v*, respectively. We enforce the consistency between these decision variables with the constraint C1. We also have to ensure that every node is visited at most once (C2) and the tour is connected and starts from and ends in *s* (C3). We then constrain the edge weight limit (C4) and the length (C5) of each tour. Additionally, we include constraints to eliminate potential subtours (i.e., solutions that include two or more disconnected tours) in C6. We are using the Miller-Tucker-Zemlin formulation [30], but any other formulation could be used as well. Constraints of the form C7 are optional and can be used to enforce a minimum overall coverage of Θ different HLA alleles or pathogens among all tours, whereas C8 can be used to enforce a minimum *average* conservation (number of antigens covered by each epitope) of Γ. C7 and C8 require a set *O* with the available options (HLA alleles or pathogen) and indicator variables *τ*_*ij*_ specifying whether epitope *i* covers option *j*. Note that C7 and/or C8 have to be repeated for every type of option, according to the requirements for the vaccine. For example, in order to enforce minimum HLA and pathogen coverage, as well as minimum average epitope conservation among pathogens, one would need *two* sets of options: *O*_*a*_for alleles and *O*_*p*_ for pathogens. Accordingly, there should be two sets of indicator variables *τ*_*va*_ and 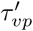 the former indicating whether *v* covers the allele *a* ∈ *O*_*a*_, and the latter indicating whether *v* covers the pathogen *p*′ ∈ *O*_*p*_. Finally, one would need to duplicate C7, one using *O*_*a*_ and the related indicators *τ*‥, and the other using *O*_*v*_ and indicators 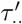, while C8 should be computed with *O*_*p*_ and 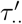.

### Data and preprocessing

#### Dataset

We downloaded 2241 sequences of the Nef gene of HIV-1 strains subtypes B and C from the Los Alamos HIV database [31, 32]. This set contained 1917 unique sequences which were used to create five bootstraps, each composed of 300 randomly selected sequences (with replacement). This was done for computational ease, as well as to study the variability and the generalizability of the results. The remaining steps of the pipeline, including the evaluation, were executed separately for each subset and the results aggregated at the end.

#### HLA alleles

We used 27 HLA alleles and their frequencies found in Toussaint *et al.* [14], reproduced in Supplement 2, which together provide a maximum theoretical coverage of 91.3% of the world population.

#### Epitopes

The binding affinities between peptides and HLA alleles were predicted with NetMHC-pan [24]. No filtering was done unless explicitly stated.

#### Cleavage likelihood

The cleavage likelihoods between all pairs of epitopes, necessary for the string-of-beads design, were predicted using PCM scores [26]. The negative cleavage likelihood of a sequence composed of two joined epitopes *e*_*i*_ and *e*_*j*_, with *e*_*i*_ of length *l*, was computed by adding to the negative cleavage score at the correct position *l* (i.e., between *e*_*i*_ and *e*_*j*_) the cleavage scores at *K* wrong positions around *l*, weighted by a factor 0 ≤ *β* ≤ 1:

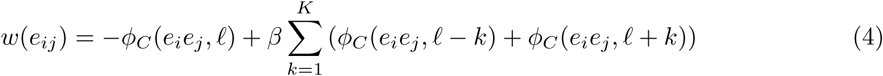

where *ϕ*_*C*_ (*s, i*) is the cleavage score at position *i* of the sequence *s*. Note that large values of *ϕ*_*C*_ (*s, i*) indicate high probability of cleavage, while large values of *w*(*e*_*ij*_) indicate high probability of incorrect cleavage. We used *K* = 2 and *β* = 0.1.

### Evaluation Metrics

Given a set of epitopes *P* comprising the vaccine, we compute the following metrics, in addition to the immunogenicity *I*(*P*):

#### Population Coverage

Given a set of HLA alleles *A*, we define the population coverage of *P* as the probability that a person has at least one HLA allele binding to one or more epitopes of *P* [14]:

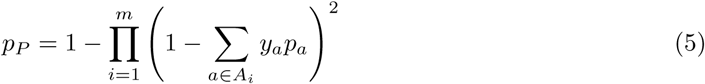

where *A*_*i*_ is the set of alleles of locus *i, p*_*a*_ is the probability that a person has this allele, and *y*_*a*_ is a binary variable indicating whether *P* contains an epitope binding to *a*. Binding was determined by an IC_50_ affinity of at most 5000 nM. Note that the graph can contain some epitopes that do not bind to any HLA allele.

We will only show the population coverage relative to the maximum that can be achieved with the given set of alleles, so that 100% relative coverage corresponds to 93.1% actual coverage.

#### Pathogen Coverage

The number of distinct pathogens that contain at least one epitope of *P*.

#### Conservation

The average of the conservation of each epitope in *P*. The conservation of an epitope is the number of proteins that contain it. High conservation indicates low mutation rate, hence importance for the correct functioning of the pathogen.

## Results

### Jointly approaching epitope selection and assembly captures trade-off between cleavage likelihood and immunogenicity

Vaccines are cleaved by the proteasome, and the resulting peptides are eventually presented on the surface of the cell by the MHC-I complex. A string-of-beads vaccine is effective only if the proteasome correctly cleaves the epitopes contained in the vaccine. Wrong cleavage sites would result in new, unwanted peptides with unknown properties, thereby decreasing the efficacy of the vaccine.

This risk can be managed with our framework by appropriately setting *h*, the maximum total negative cleavage log-likelihood between all epitopes of the vaccine. Interpreting and predetermining this quantity is, however, difficult. For each of the five bootstraps, we generated several Pareto-efficient solutions to illustrate the trade-off between correct cleavage and high immunogenicity.

Cleavage likelihood can be increased considerably with small losses in immunogenicity up until a certain point, after which the latter quickly drops to provide only modest improvements in cleavage (Figure 2). Notably, it is possible to achieve 84-88% of the maximum cleavage score with only a 15-20% reduction in immunogenicity. In practice, the effective immunogenicity for vaccines with low cleavage likelihood is smaller than the theoretical immunogenicity, as the vaccine is more likely to be cleaved incorrectly. Consequently, the cleavage likelihood should be favored more than immunogenicity when choosing the design parameters.

**Figure 2:**
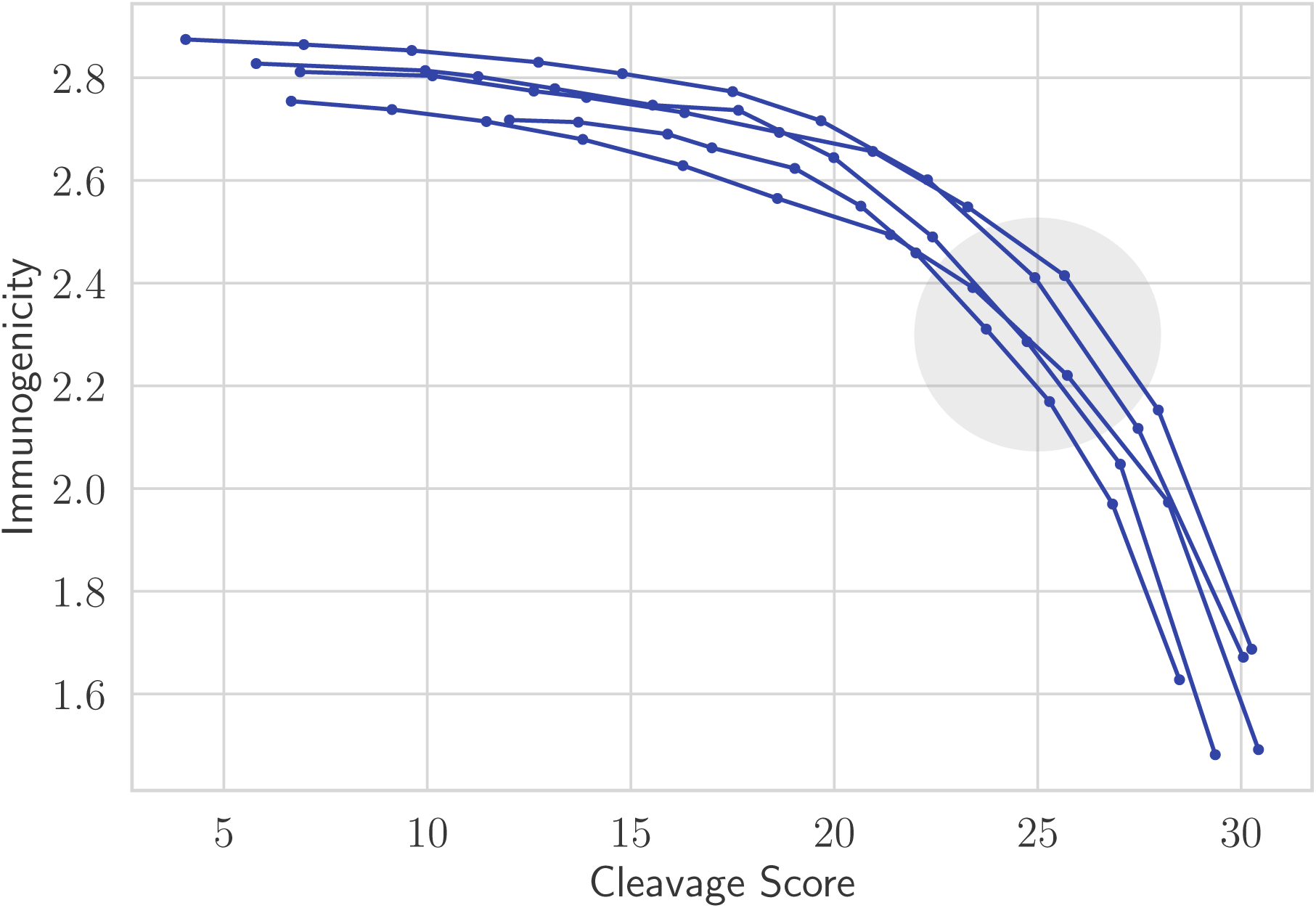
Novel EV design possibilities opened by considering epitope selection and epitope assembly at the same time. Pareto frontiers between immunogenicity and cleavage score for five different bootstraps show that it is possible to obtain a vaccine that is processed reasonably well by the proteasome without compromising its immunogenicity. The color encodes the local slope, and the circle identifies the region that should be preferred when designing vaccines.

Sequential approaches such as OptiTope [13] are only able to generate the highest-immunogenicity solutions, as they do not consider the subsequent assembly phase as all. These methods simply cannot be used to balance the quality of selection and assembly.

### Joint design of polypeptide cocktails increases immunogenicity without sacrifices

A large number of epitopes might be necessary to create a vaccine meeting extreme requirements. Long polypeptides, however, are harder to manufacture [33, 34] and, in practice, most synthetic vaccines tested so far are composed of sequences of 10 to 50 amino acids [4, 35–38]. Our framework can be used to design a vaccine that meets extreme requirements with short polypeptides that are optimized simultaneously, and can be synthesized in parallel for a fraction of the cost and time.

An epitope mixture designed by OptiTope [13] needs at least 24 epitopes to cover 99% of the complete set of pathogens, but 216 amino acids might be too many for a single string-of-beads polypeptide. We, therefore, designed a vaccine composed of four separate mosaic polypeptides of 54 amino acids each. We also designed a single mosaic of 216 amino acids with no coverage enforced as a baseline. The mosaic cocktail was designed by considering the union of the 2000 epitopes with highest immunogenicity and the 2000 epitopes with highest pathogen coverage.

The resulting four polypeptides together have roughly twice the conservation and the immunogenicity as the epitope mixture (Figure 3). Most notably, none of them reach the required pathogen coverage in isolation; only when considered jointly they cover the required number of pathogens. The unconstrained single mosaic vaccine already covers 96% of the pathogens even though its epitopes have the lowest conservation among the three vaccines. Unsurprisingly, it also has the largest immunogenicity, almost 20% more than the polypeptide cocktail and 260% more than the epitope mixture.

**Figure 3:**
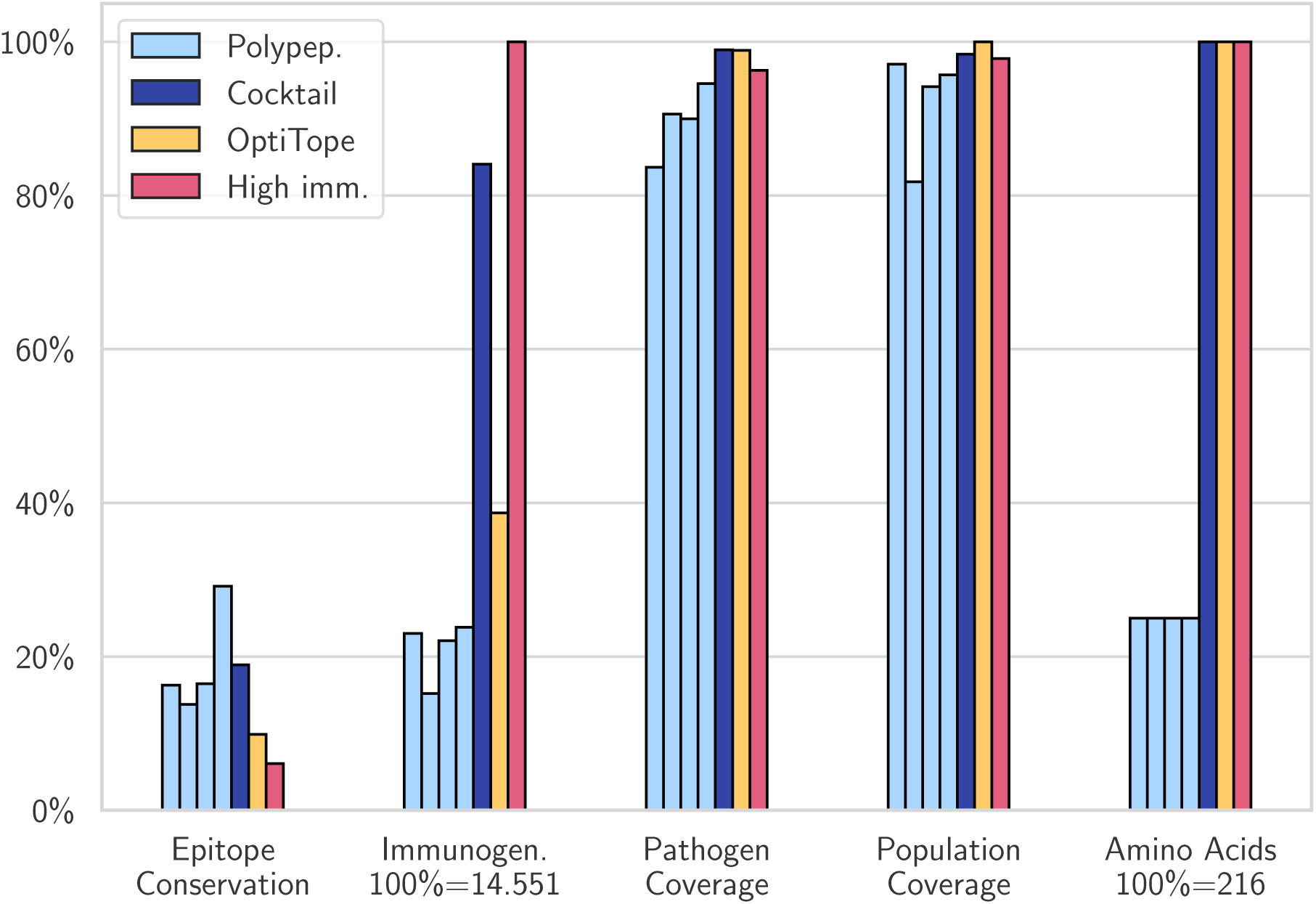
We designed a cocktail (white) of four polypeptides (cyan) that covers 99% of the pathogens, even though the single fragments only cover between 85 and 95%. The orange and red columns correspond to an epitope mixture designed by OptiTope and a mosaic with the same number of amino acids respectively; the former was required to reach 99% pathogen coverage, and the latter was unconstrained.

### Mosaic design greatly increases immunogenicity and pathogen coverage compared to string-of-beads

By leveraging overlaps, the mosaic design is able to include more epitopes in the same number of amino acids, resulting in improved immunogenicity and coverage compared to the string-of-beads design (Figure 4). This occurs as the enforced overlap cannot be sustained by the limited variety of the input epitopes. By relaxing this constraint to only four amino acids, we can produce pseudo-mosaic vaccines that contain less epitopes than the theoretical maximum, but have, nonetheless, much higher immunogenicity than the string-of-beads alternative. It is also evident that mosaic vaccines can inherently reach higher pathogen coverage with shorter polypeptides, even when no such constraint is imposed on the design. Conservation, however, remains mediocre.

**Figure 4:**
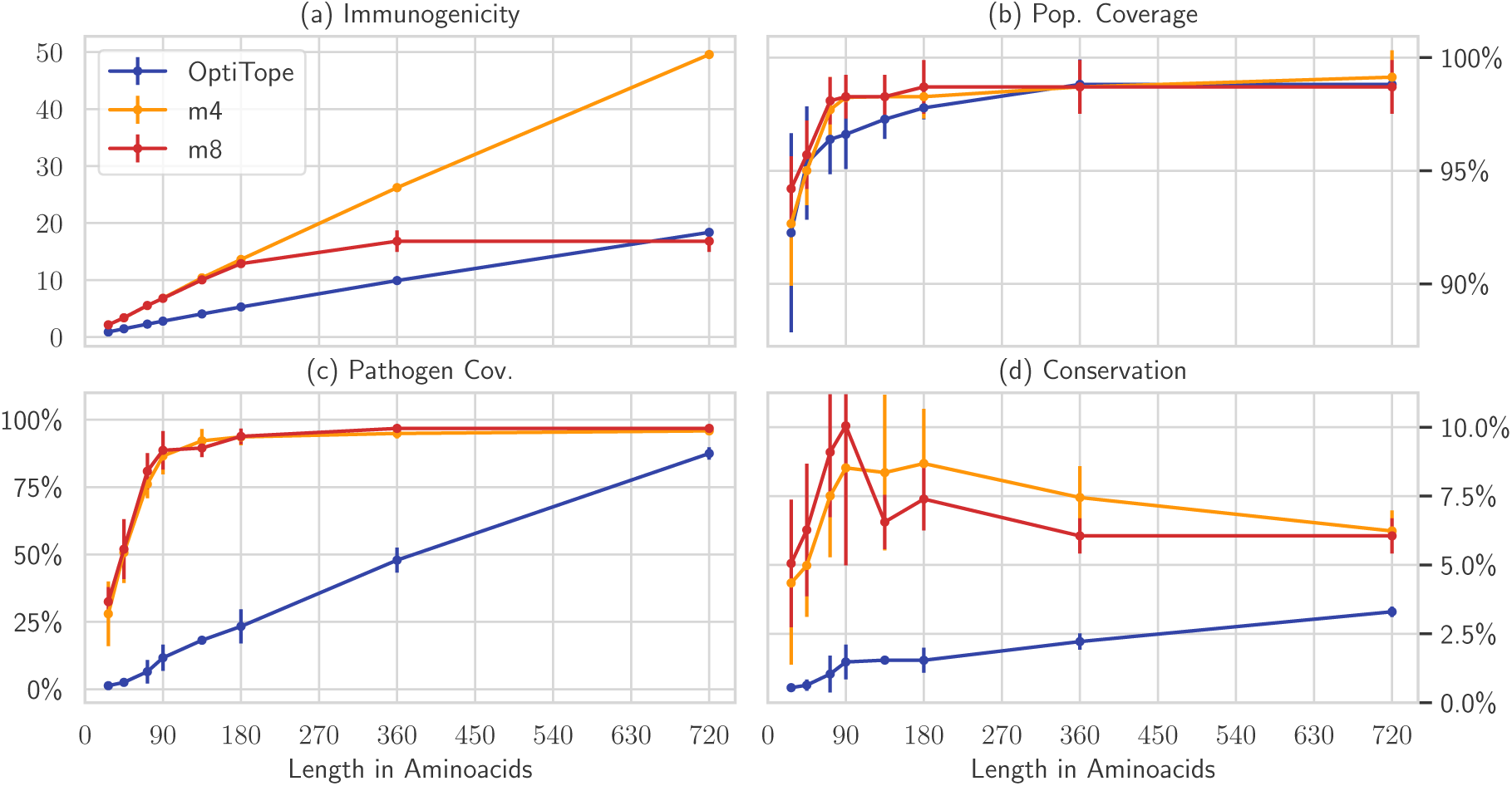
Mosaic vaccines are much better than epitope mixtures or string-of-beads of the same length designed by OptiTope (blue), as long as the pathogens offer enough epitope variety. By enforcing an overlap between epitopes of eight amino acids (red), the vaccine does not improve after a certain length. This can be prevented by relaxing this requirement to only four amino acids (yellow). The vaccines are compared with respect to four metrics: immunogenicity (a), population coverage (b), pathogen coverage(c), and conservation (d). Bars represent the standard deviation of five separate runs. Note that these vaccines were not designed with pathogen coverage nor epitope conservation in mind; mosaics are naturally better at this.

### Short mosaic vaccines achieve very high coverage

We designed mosaic vaccines with a single polypeptide on each of the five bootstraps following the genetic algorithm introduced by Fisher *et al.* [19, 39], using the recommended parameter settings (notably, unlike our framework, the length of the polypeptides cannot be specified). We then used our framework to design a mosaic vaccine of the same length (206 amino acids), with at least the same pathogen and population coverage and epitope conservation. This results in very similar mosaics with essentially the same properties: the same 26 HLA alleles covered out of 27, 99.1% of pathogens covered, an average epitope conservation of 28%, and immunogenicity of 10.8. However, Fisher *et al.* optimize for coverage, not for immunogenicity. If we do the same, we are able to achieve the same properties with a reduction in epitopes of about 50%.

### Long mosaic vaccines inherently target conserved regions

The previous experiments show that mosaic vaccines can reach very good pathogen coverage with ease. Intrigued by this characteristic, we studied and compared the exact positions covered by the epitopes of mosaic and string-of-beads vaccines.

The vaccines were designed on the complete pathogen set. The string-of-beads contained 20 epitopes, the short mosaic vaccine 28 amino acids, and the long mosaic vaccine 90 amino acids. We then aligned the sequences using MAFFT [40] and counted how many epitopes cover every position. We also computed the potential immunogenicity as the sum of the immunogenicities of all the epitopes covering that position, and used position-specific entropy to quantify the variation among sequences (section B in Supplement 1). Finally, we ignored the positions where the consensus (majority) was a gap.

Analyzing the epitopes included in the vaccines showed that they do not appear in random positions of the pathogens, but are concentrated in a few distinct regions that differ between vaccines (Figure 5 B). It is evident that string-of-beads vaccines, cleavage requirements aside, target the most immunogenic regions with no regards for their conservation, whereas mosaic vaccines, especially longer ones, prefer to focus on conserved regions. These correlations, as quantified by the Spearman coefficient, are generally weak or moderate, but statistically significant (Figure 5 C). The epitope mixture’s coverage was well correlated with immunogenicity (*r* = 0.617, p=4 · 10^-22^), but not with entropy (*r* = 0.066, *p* = 4 · 10^-1^). The long mosaic vaccine seeked immunogenic (*r* = 0.350, *p* = 4·10^-7^), but low entropy regions (*r* = −0.353, *p* = 3·10^-7^). Interestingly, the short mosaic vaccine covered an entirely different region and was correlated with both immunogenicity (*r* = 0.228, *p* = 1 · 10^-3^) and entropy (*r* = 0.410, 2 · 10^-9^). Entropy and immunogenicity are, curiously, not correlated (*r* = −0.015, *p* = 8 · 10^-1^).

**Figure 5:**
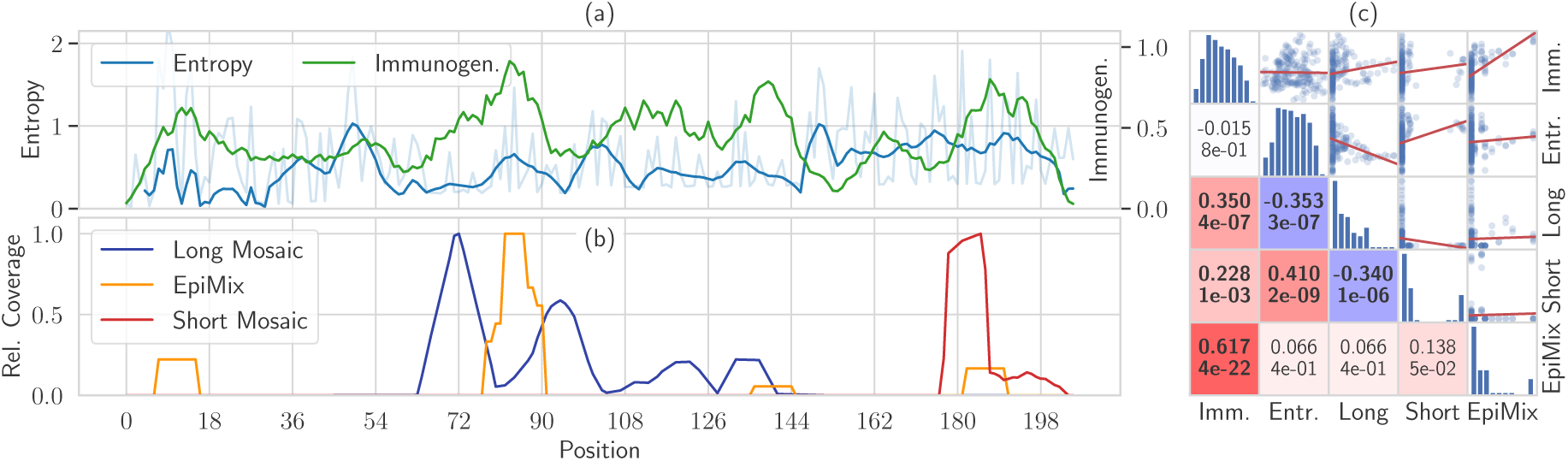
Mosaics vaccines naturally target conserved regions even when this is not required. **(a):** shows, for each residue position in aligned sequences where the consensus is not a gap, the smoothed entropy (blue, and residue entropy in lighter color) and the potential immunogenicity (green) **(b):** shows the number of pathogens covered in each position by a 20-epitopes mixture with maximal immunogenicity (yellow), a short mosaic of 28 amino acids (red) and a long mosaic of 0 amino acids (blue). The count is normalized separately for each vaccine to account for their different coverage. **(c):** shows the pairwise correlations of the variables shown in the left plot, so that every dot in the scatter plots corresponds to a different residue position, and linear fits are shown in red. The lower triangular half shows the Spearman correlation coefficients (above) and the respective *p*-value (below). Colors range from blue (large negative correlation) to white (no correlation) to red (large positive correlation), and the font is bold if the correlation is significant with a confidence of at least 99.5%, the Bonferroni-corrected standard significance level of 5%. The diagonal contains histograms showing the distribution of each variable, with logarithmic *y* axis.

### Mosaic vaccines should be designed with epitope conservation in mind

There is growing evidence that effective vaccines for highly variable viruses such as the Human Immunodeficiency Virus (HIV), Hepatitis C Virus (HCV), as well as diseases such as Malaria, Cancer, and Influenza should target conserved epitopes [41–45]. Some of our previous experiments clearly showed that mosaic vaccines have a natural tendency to spontaneously achieve high pathogen coverage, even though the conservation of the individual epitopes in the vaccine remains low.

We modified the ILP formulation to maximize average epitope conservation and pathogen coverage together with immunogenicity (Supplement 3). We then compared mosaic vaccines of growing sizes optimized against these three criteria (Figure 6). Since immunogenicity is a couple of orders of magnitude smaller than the other two, it will only be optimized when further improvements in conservation or coverage are practically insignificant. The average epitope conservation can be greatly improved until around 40%, and it comes with increased pathogen coverage compared to mosaic vaccines optimized for immunogenicity. Most epitopes have very poor conservation, which means that optimizing its average becomes harder as the vaccine size increases. In fact, conservation decreases for longer vaccines. Moreover, immunogenicity grows more slowly when conservation is optimized: it is on par with pathogen coverage-optimized vaccines for short mosaic designs, but is almost 30% smaller for long mosaic designs.

**Figure 6:**
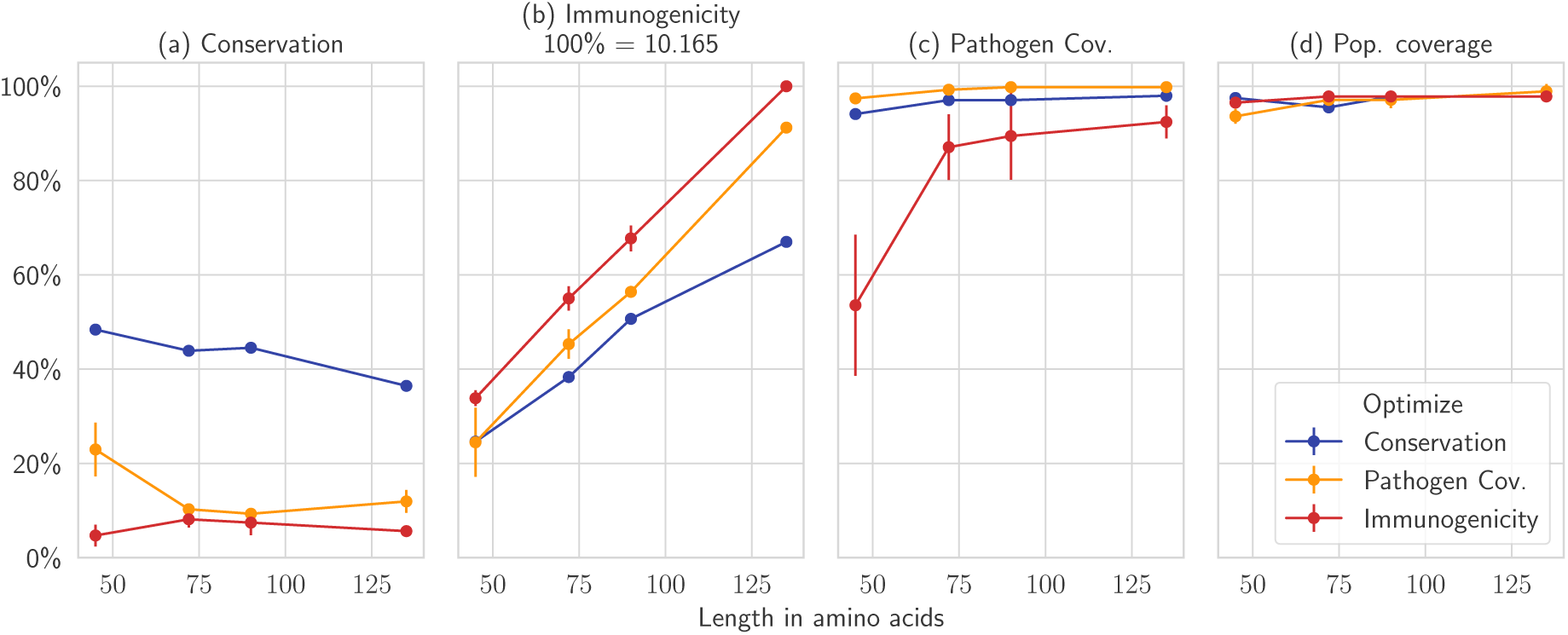
Here we design mosaics of varying size (on the *x* axis) while optimizing for conservation (blue), immunogenicity (red) and pathogen coverage (yellow). The plots compare the vaccines in terms of conservation (a), immunogenicity (b), pathogen coverage (c), and population coverage (d). For longer vaccines, optimizing for pathogen coverage only gives modest improvements on the mosaics optimized for immunogenicity in terms of coverage, and does not increase conservation by much. When optimized for, conservation is considerably higher, but becomes harder to improve as the vaccine becomes longer, due to the fact that few epitopes are well conserved and highly immunogenic at the same time. Average of five runs, standard deviation on the error bars.

This suggests that the best results are obtained with short mosaic vaccines designed to have high average epitope conservation. Besides having considerably larger conservation, both their immunogenicity and their pathogen coverage are still close to the theoretical maximum. As the mosaic designs become longer, this gap widens, and so-designed vaccines lose their advantages. However, we have shown previously that long vaccines can be replaced by cocktails of short polypeptides with essentially the same joint properties.

### Vaccines designed on small subsets generalize to the full dataset

Some of the previous results were obtained by only considering 300 random proteins out of a few thousand, and the cocktail only used a subset of about 4000 out of 52712 epitopes. One naturally wonders whether the produced vaccines are as good on the general pathogen population.

The only quantities that can change, for a given vaccine, are epitope conservation and pathogen coverage, while designing a vaccine *de novo* can result in higher immunogenicity. We use the same setting of the experiment where we compared with Fischer *et al* and consider it representative of the others, since we mostly focused our attention to mosaics. We evaluated the mosaics designed on the five bootstrap on the set of all pathogens, and quantified the differences in the evaluation metrics. This revealed that there was no difference on conservation (paired t-test, *t* = 0.04, *p* = 0.487) between the small subsets and the full set, but pathogen coverage slightly decreased both for our design and for Fischer *et al.*’s (*t* = 2.77, *p* = 0.025, from 99.1%, std. 0.4% to 98.7%, std. 0.2%; similar quantities for [19]). As for immunogenicity, a string-of-beads with 10 epitopes and no constraints achieved an immunogenicity of 3.08 on the full set, just 10% higher than what could be achieved with the same settings on the smaller sets in the Pareto frontier experiment. A 216-amino acids mosaic in the same setting of the cocktail experiment, but designed on all the epitopes, improved immunogenicity by 18% (from 14.55 to 17.23), but worsened coverage by 1% (from 96.3% to 95.0%) and conservation by 41% (from 6.1% to 3.6%), suggesting that subsetting epitopes should be done with greater care.

It might seem surprising that vaccines developed on roughly 15% of the pathogens generalize to a larger population. However, peptides with high coverage and conservation on a large population are likely to be so also on random subsets of it. In fact, even though each of the five random subsets contains only about 25% of the peptides found in the full pathogen set, the pairwise overlap is between 46 and 48%, and 27% of their peptides are shared among all five sets.

## Discussion and conclusion

Epitope-based vaccine (EV) design has thriven in recent years, and multiple design principle have emerged aided by the heavy use of bioinformatics approaches. However, most proposed design algorithms are lacking in one of several dimensions: they model only individual stages of the entire design problem (e.g., [13, 15]), use *ad hoc* heuristics (e.g., [16]), or optimization algorithms that cannot guarantee convergence to the optimal solution (e.g., [12, 19]).

Here we proposed a graph-theoretical formalism for EV design that models the complete design process and includes every prevalent design principle as special case. We showed how to formulate this optimization problem as an integer linear program to obtain an guaranteed optimal solution. This, in turn, enables informed choices throughout the design process by accurately and reliably determining the trade-offs involved: for example, we precisely quantified the decrease in immunogenicity that has to be paid to achieve gains in other metrics such as coverage and conservation, and showed the advantage of mosaic over string-of-beads designs under our modeling assumptions. In practice, we might be overestimating mosaics’ immunogenicity, as our framework does not model their cleavage by the proteasome, which means that we have no control over which epitopes will actually be recovered. Their successes in recent clinical trials [5–10, 17], however, suggest that their advantage over string-of-beads is, nonetheless, real. Jointly approaching the selection and assembly problems enables the exploration of new possibilities in the EV design space. We demonstrated this by investigating the trade-off between immunogenicity and cleavage likelihood in string-of-beads designs, and by creating the optimal cocktail of mosaic polypeptides for a given coverage whose design would be impossible with iterative, stage-wise optimization methods. Our results also show that it is easy to reach very good population coverage by virtue of our definition of immunogenicity based on HLA binding. Finally, convergence and optimality guarantees of linear programming solvers allow us to find solutions that are, sometimes, substantially better than those found by optimization algorithms that lack these guarantees.

The price to pay for the increased modeling power is increased computational resources needed to solve the graph optimization problem. Being based on the team orienteering problem, EV design with our framework is a NP-hard problem, and the size of the graph grows quadratically with the number of epitopes in consideration. However, we conducted several experiments on subsets of the pathogens or the epitopes, and showed that results obtained in this way are only slightly worse than what can be obtained by considering the complete set of pathogens/epitopes. We argued that this is possible because highly conserved epitopes are likely to abound in smaller subsets of the pathogen sequences too. This means that such graphs can easily be pruned, resulting in much smaller problems that can be solved in reasonable time without compromising the quality of the final solution. Alternatively, the solver can be interrupted early when the current solution is within a few percents of the optimal one. ILP solvers iteratively improve a candidate solution and an upper bound on the objective of the optimal solution at the same time [46]. This gap tells us the maximum distance between the candidate solution and the optimal one: when they match, the current solution is optimal. Empirically, in our setting, most of the time is spent on improving solutions that are only at most 2%-5% away from the optimal one. Therefore, the solver can be safely interrupted early on a good quality solution. ILP solvers are complex machinery governed by several parameters that affect how they search for a solution, and, therefore, how much time they need. Different types of problems benefit from different parameter settings. Therefore, further gains can be achieved when dealing with a large number of instances by tuning these parameters to reduce the time needed to find a solution [47–49]. Another limitation is that our formulation as an ILP limits the expressiveness of objectives and constraints to linear forms. The graph formalism in Eq. 1, however, remains valid even for complex, non-linear constraints and objectives. In this case, more flexible optimization methods have to be used, but the optimality guarantees can be lost.

To conclude, the proposed framework enables the design of correctly cleaved string-of-beads vaccines with the largest possible immunogenicity under this constraint. It also enables the exploration of the Pareto frontier between these two competing properties to find the optimal design that best reflects the envisioned trade-off. At the same time, our framework can be used to decompose long vaccines in shorter polypeptides that, together, maintain the properties of the longer sequence. This makes the resulting vaccine easier, cheaper, and quicker to synthesize. Finally, we showed that conservation should be emphasized over coverage for mosaic vaccines.

## Supporting information

Supplement One

Supplement Two

Supplement Three

## Declarations

### Availability of data and materials

The code, as well as any datasets generated and/or analysed during the study is available at https://github.com/SchubertLab/GeneralizedEvDesign

### Authors’ contributions

Both authors developed the mathematical framework. ED implemented the software, performed the experiments and statistical analyses. BS designed the study. Both authors wrote, read, and approved the final manuscript.

## Acknowledgments

We would like to express our gratitude to Dr. Bernd Bischl for his comments on the manuscript.

## Supplementary material

**Supplement 1:** (PDF file) section A contains a brief description of the *E*-constrain method [28] to obtain Pareto-efficient solutions in a bi-objective optimization problem, while section B contains the procedure to quantify conserved, low variability pathogen regions in terms of entropy on aligned sequences.

**Supplement 2:** (PDF file) contains a table listing the 27 HLA alleles used in this study and their percent frequency in the world population.

**Supplement 3:** (PDF file) contains the two alternative formulations of the ILP where average epitope conservation and pathogen coverage are maximized together with immunogenicity.

## References

[1] J. C. Castle, S. Kreiter, J. Diekmann, M. Lower, N. van de Roemer, J. de Graaf, A. Selmi, M. Diken, S. Boegel, C. Paret, M. Koslowski, A. N. Kuhn, C. M. Britten, C. Huber, O. Tureci, and U. Sahin. Exploiting the Mutanome for Tumor Vaccination. Cancer Research, 72(5):1081–1091, March 2012.

[2] Sebastian Kreiter, Mathias Vormehr, Niels van de Roemer, Mustafa Diken, Martin Löwer, Jan Diekmann, Sebastian Boegel, Barbara Schrörs, Fulvia Vascotto, John C. Castle, Arbel D. Tadmor, Stephen P. Schoenberger, Christoph Huber, Ö zlem Türeci, and Ugur Sahin. Mutant MHC class II epitopes drive therapeutic immune responses to cancer. Nature, 520(7549):692–696, April 2015.

[3] Hirokazu Matsushita, Matthew D. Vesely, Daniel C. Koboldt, Charles G. Rickert, Ravindra Uppaluri, Vincent J. Magrini, Cora D. Arthur, J. Michael White, Yee-Shiuan Chen, Lauren K. Shea, Jasreet Hundal, Michael C. Wendl, Ryan Demeter, Todd Wylie, James P. Allison, Mark J. Smyth, Lloyd J. Old, Elaine R. Mardis, and Robert D. Schreiber. Cancer exome analysis reveals a T-cell-dependent mechanism of cancer immunoediting. Nature, 482(7385):400–404, February 2012.

[4] Patrick A. Ott, Zhuting Hu, Derin B. Keskin, Sachet A. Shukla, Jing Sun, David J. Bozym, Wandi Zhang, Adrienne Luoma, Anita Giobbie-Hurder, Lauren Peter, Christina Chen, Oriol Olive, Todd A. Carter, Shuqiang Li, David J. Lieb, Thomas Eisenhaure, Evisa Gjini, Jonathan Stevens, William J. Lane, Indu Javeri, Kaliappanadar Nellaiappan, Andres M. Salazar, Heather Daley, Michael Seaman, Elizabeth I. Buchbinder, Charles H. Yoon, Maegan Harden, Niall Lennon, Stacey Gabriel, Scott J. Rodig, Dan H. Barouch, Jon C. Aster, Gad Getz, Kai Wucherpfennig, Donna Neuberg, Jerome Ritz, Eric S. Lander, Edward F. Fritsch, Nir Hacohen, and Catherine J. Wu. An immunogenic personal neoantigen vaccine for patients with melanoma. Nature, 547(7662):217–221, July 2017.

[5] Dan H Barouch, Kara L O’Brien, Nathaniel L Simmons, Sharon L King, Peter Abbink, Lori F Maxfield, Ying-Hua Sun, Annalena La Porte, Ambryice M Riggs, Diana M Lynch, Sarah L Clark, Katherine Backus, James R Perry, Michael S Seaman, Angela Carville, Keith G Mansfield, James J Szinger, Will Fischer, Mark Muldoon, and Bette Korber. Mosaic HIV-1 vaccines expand the breadth and depth of cellular immune responses in rhesus monkeys. Nature Medicine, 16(3):319–323, March 2010.

[6] W.-P. Kong, L. Wu, T. C. Wallstrom, W. Fischer, Z.-Y. Yang, S.-Y. Ko, N. L. Letvin, B. F. Haynes, B. H. Hahn, B. Korber, and G. J. Nabel. Expanded Breadth of the T-Cell Response to Mosaic Human Immunodeficiency Virus Type 1 Envelope DNA Vaccination. Journal of Virology, 83(5):2201–2215, March 2009.

[7] Sampa Santra, Mark Muldoon, Sydeaka Watson, Adam Buzby, Harikrishnan Balachandran, Kevin R. Carlson, Linh Mach, Wing-Pui Kong, Krisha McKee, Zhi-Yong Yang, Srinivas S. Rao, John R. Mascola, Gary J. Nabel, Bette T. Korber, and Norman L. Letvin. Breadth of cellular and humoral immune responses elicited in rhesus monkeys by multi-valent mosaic and consensus immunogens. Virology, 428(2):121–127, July 2012.

[8] Dan H. Barouch, Kathryn E. Stephenson, Erica N. Borducchi, Kaitlin Smith, Kelly Stanley, Anna G. McNally, Jinyan Liu, Peter Abbink, Lori F. Maxfield, Michael S. Seaman, Anne-Sophie Dugast, Galit Alter, Melissa Ferguson, Wenjun Li, Patricia L. Earl, Bernard Moss, Elena E. Giorgi, James J. Szinger, Leigh Anne Eller, Erik A. Billings, Mangala Rao, Sodsai Tovanabutra, Eric Sanders-Buell, Mo Weijtens, Maria G. Pau, Hanneke Schuitemaker, Merlin L. Robb, Jerome H. Kim, Bette T. Korber, and Nelson L. Michael. Protective Efficacy of a Global HIV-1 Mosaic Vaccine against Heterologous SHIV Challenges in Rhesus Monkeys. Cell, 155(3):531–539, October 2013.

[9] Dan H. Barouch, Frank L. Tomaka, Frank Wegmann, Daniel J. Stieh, Galit Alter, Merlin L. Robb, Nelson L. Michael, Lauren Peter, Joseph P. Nkolola, Erica N. Borducchi, Abishek Chandrashekar, David Jetton, Kathryn E. Stephenson, Wenjun Li, Bette Korber, Georgia D. Tomaras, David C. Montefiori, Glenda Gray, Nicole Frahm, M. Juliana McElrath, Lindsey Baden, Jennifer Johnson, Julia Hutter, Edith Swann, Etienne Karita, Hannah Kibuuka, Juliet Mpendo, Nigel Garrett, Kathy Mngadi, Kundai Chinyenze, Frances Priddy, Erica Lazarus, Fatima Laher, Sorachai Nitayapan, Punnee Pittisuttithum, Stephan Bart, Thomas Campbell, Robert Feldman, Gregg Lucksinger, Caroline Borremans, Katleen Callewaert, Raphaele Roten, Jerald Sadoff, Lorenz Scheppler, Mo Weijtens, Karin Feddes-de Boer, Daniëlle van Manen, Jessica Vreugdenhil, Roland Zahn, Ludo Lavreys, Steven Nijs, Jeroen Tolboom, Jenny Hendriks, Zelda Euler, Maria G. Pau, and Hanneke Schuitemaker. Evaluation of a Mosaic HIV-1 Vaccine in a Randomized, Double-Blinded, Placebo-Controlled Phase I/IIa Clinical Trial and in Rhesus Monkeys. Lancet (London, England), 392(10143):232–243, July 2018.

[10] Lindsey R Baden, Etienne Karita, Gaudensia Mutua, Linda-Gail Bekker, Glenda Gray, Liesl Page-Shipp, Stephen R Walsh, Julien Nyombayire, Omu Anzala, Surita Roux, Fatima Laher, Craig Innes, Michael Seaman, Yehuda Z Cohen, Lauren Peter, Nicole Frahm, M Juliana McElrath, Peter Hayes, Edith Swann, Nicole Grunenberg, Maria Grazia-Pau, Mo Weijtens, Jerry Sadoff, Len Dally, Angela Lombardo, Jill Gilmour, Josephine Cox, Raphael Dolin, Patricia Fast, Dan H Barouch, Dagna S Laufer, Jennifer Johnson, Jane Kleinjan, Rosine Ingabire, Delvin Nyasani, Danielle Crida, Nicholas Mangeya, Musawenkosi Mamba, Kathy Mngadi, David J Dominguez, Katherine E Yanosick, Emmanuel Cormier, John Hural, Gwynn Stevens, Elizabeth Adams, James Kublin, Jenny Hendriks, Eddy Sayeed, James Ackland, Kamaal Anas, Devika Zackariah, Dani Vooijs, Kundai Chinyenze, Mabela Matsoso, Harriet Park, and Sabrina Welsh. Assessment of the Safety and Immunogenicity of 2 Novel Vaccine Platforms for HIV-1 Prevention: A Randomized Trial. Annals of internal medicine, 164(5):313–322, March 2016.

[11] Anthony W. Purcell, James McCluskey, and Jamie Rossjohn. More than one reason to rethink the use of peptides in vaccine design. Nature Reviews Drug Discovery, 6(5):404–414, May 2007.

[12] Tal Vider-Shalit, Shai Raffaeli, and Yoram Louzoun. Virus-epitope vaccine design: Informatic matching the HLA-I polymorphism to the virus genome. Molecular Immunology, 44(6):1253–1261, February 2007.

[13] Nora C. Toussaint, Pierre Dönnes, and Oliver Kohlbacher. A Mathematical Framework for the Selection of an Optimal Set of Peptides for Epitope-Based Vaccines. PLoS Computational Biology, 4(12):e1000246, December 2008.

[14] Nora C. Toussaint, Yaakov Maman, Oliver Kohlbacher, and Yoram Louzoun. Universal peptide vaccines – Optimal peptide vaccine design based on viral sequence conservation. Vaccine, 29(47):8745–8753, November 2011.

[15] Benjamin Schubert and Oliver Kohlbacher. Designing string-of-beads vaccines with optimal spacers. Genome Medicine, 8(1), December 2016.

[16] C. Lundegaard, M. Buggert, Ac Karlsson, O. Lund, Carina Perez, and M. Nielsen. PopCover: a method for selecting of peptides with optimal population and pathogen coverage. In Proceedings of the First ACM International Conference on Bioinformatics and Computational Biology - BCB ‘10, page 658, Niagara Falls, New York, 2010. ACM Press.

[17] Sultan Abdul-Jawad, Beatrice Ondondo, Andy van Hateren, Andrew Gardner, Tim Elliott, Bette Korber, and Tomáš Hanke. Increased Valency of Conserved-mosaic Vaccines Enhances the Breadth and Depth of Epitope Recognition. Molecular Therapy, 24(2):375–384, February 2016.

[18] Tracey A Day, Cecilia A Morgan, and James G Kublin. Will mosaic vaccine immunogens expand immune response breadth to rival HIV-1 strain diversity? Clinical Investigation, 3(5):413–415, May 2013.

[19] Will Fischer, Simon Perkins, James Theiler, Tanmoy Bhattacharya, Karina Yusim, Robert Funkhouser, Carla Kuiken, Barton Haynes, Norman L Letvin, Bruce D Walker, Beatrice H Hahn, and Bette T Korber. Polyvalent vaccines for optimal coverage of potential T-cell epitopes in global HIV-1 variants. Nature Medicine, 13(1):100–106, January 2007.

[20] Sébastien Cornet, Isabelle Miconnet, Jeanne Menez, François Lemonnier, and Kostas Kosmatopoulos. Optimal organization of a polypeptide-based candidate cancer vaccine composed of cryptic tumor peptides with enhanced immunogenicity. Vaccine, 24(12):2102–2109, 2006.

[21] Marco Caserta and Stefan Voß. A hybrid algorithm for the DNA sequencing problem. Discrete Applied Mathematics, 163:87–99, January 2014.

[22] Pieter Vansteenwegen, Wouter Souffriau, and Dirk Van Oudheusden. The orienteering problem: A survey. European Journal of Operational Research, 209(1):1–10, February 2011.

[23] Bruce L. Golden, Larry Levy, and Rakesh Vohra. The orienteering problem. Naval Research Logistics, 34(3):307–318, June 1987.

[24] Vanessa Isabell Jurtz, Sinu Paul, Massimo Andreatta, Paolo Marcatili, Bjoern Peters, and Morten Nielsen. NetMHCpan 4.0: Improved peptide-MHC class I interaction predictions integrating eluted ligand and peptide binding affinity data. bioRxiv, September 2017.

[25] S. Tenzer, B. Peters, S. Bulik, O. Schoor, C. Lemmel, M. M. Schatz, P.-M. Kloetzel, H.-G. Rammensee, H. Schild, and H.-G. Holzhütter. Modeling the MHC class I pathway by combining predictions of proteasomal cleavage,TAP transport and MHC class I binding. CMLS Cellular and Molecular Life Sciences, 62(9):1025–1037, May 2005.

[26] Pierre Dönnes and Oliver Kohlbacher. Integrated modeling of the major events in the MHC class I antigen processing pathway. Protein Science, 14(8):2132–2140, August 2005.

[27] Morten Nielsen, Claus Lundegaard, Ole Lund, and Can Keşmir. The role of the proteasome in generating cytotoxic T-cell epitopes: insights obtained from improved predictions of proteasomal cleavage. Immunogenetics, 57(1-2):33–41, April 2005.

[28] George Mavrotas. Effective implementation of the *ϵ*-constraint method in Multi-Objective Mathematical Programming problems. Applied Mathematics and Computation, 213(2):455–465, July 2009.

[29] Dan Gusfield, Gad M. Landau, and Baruch Schieber. An Efficient Algorithm for the All Pairs Suffix-Prefix Problem. In Renato Capocelli, Alfredo De Santis, and Ugo Vaccaro, editors, Sequences II, pages 218–224. Springer New York, New York, NY, 1993.

[30] C. E. Miller, A. W. Tucker, and R. A. Zemlin. Integer Programming Formulation of Traveling Salesman Problems. Journal of the ACM, 7(4):326–329, October 1960.

[31] Brian Thomas Foley, Bette Tina Marie Korber, Thomas Kenneth Leitner, Cristian Apetrei, Beatrice Hahn, Ilene Mizrachi, James Mullins, Andrew Rambaut, and Steven Wolinsky. HIV Sequence Compendium 2018. Technical Report LA-UR-18-25673, Los Alamos National Lab. (LANL), Los Alamos, NM (United States), June 2018.

[32] Los Alamos National Laboratory. The hiv sequence database. https://www.hiv.lanl.gov. Accessed: 2019-10-03.

[33] Giampietro Corradin, Andrey V. Kajava, and Antonio Verdini. Long Synthetic Peptides for the Production of Vaccines and Drugs: A Technological Platform Coming of Age. Science Translational Medicine, 2(50):50rv3–50rv3, September 2010.

[34] Stephen B. H. Kent. Total chemical synthesis of proteins. Chemical Society Reviews, 38(2):338–351, January 2009.

[35] Martijn S. Bijker, Susan J. F. van den Eeden, Kees L. Franken, Cornelis J. M. Melief, Rienk Offringa, and Sjoerd H. van der Burg. CD8+ CTL priming by exact peptide epitopes in incomplete Freund’s adjuvant induces a vanishing CTL response, whereas long peptides induce sustained CTL reactivity. Journal of Immunology (Baltimore, Md.: 1950), 179(8):5033–5040, October 2007.

[36] Haniyeh Ghaffari-Nazari, Jalil Tavakkol-Afshari, Mahmoud Reza Jaafari, Sahar Tahaghoghi-Hajghorbani, Elham Masoumi, and Seyed Amir Jalali. Improving Multi-Epitope Long Peptide Vaccine Potency by Using a Strategy that Enhances CD4+ T Help in BALB/c Mice. PLOS ONE, 10(11):e0142563, November 2015.

[37] Ugur Sahin, Evelyna Derhovanessian, Matthias Miller, Björn-Philipp Kloke, Petra Simon, Martin Löwer, Valesca Bukur, Arbel D. Tadmor, Ulrich Luxemburger, Barbara Schrörs, Tana Omokoko, Mathias Vormehr, Christian Albrecht, Anna Paruzynski, Andreas N. Kuhn, Janina Buck, Sandra Heesch, Katharina H. Schreeb, Felicitas Müller, Inga Ortseifer, Isabel Vogler, Eva Godehardt, Sebastian Attig, Richard Rae, Andrea Breitkreuz, Claudia Tolliver, Martin Suchan, Goran Martic, Alexander Hohberger, Patrick Sorn, Jan Diekmann, Janko Ciesla, Olga Waksmann, Alexandra-Kemmer Brück, Meike Witt, Martina Zillgen, Andree Rothermel, Barbara Kasemann, David Langer, Stefanie Bolte, Mustafa Diken, Sebastian Kreiter, Romina Nemecek, Christoffer Gebhardt, Stephan Grabbe, Christoph Höller, Jochen Utikal, Christoph Huber, Carmen Loquai, and Ö zlem Türeci. Personalized RNA mutanome vaccines mobilize poly-specific therapeutic immunity against cancer. Nature, 547(7662):222–226, July 2017.

[38] S. Zwaveling, S. C. Ferreira Mota, J. Nouta, M. Johnson, G. B. Lipford, R. Offringa, S. H. van der Burg, and C. J. M. Melief. Established Human Papillomavirus Type 16-Expressing Tumors Are Effectively Eradicated Following Vaccination with Long Peptides. The Journal of Immunology, 169(1):350–358, 2002.

[39] James Thurmond, Hyejin Yoon, Carla Kuiken, Karina Yusim, Simon Perkins, James Theiler, Tanmoy Bhattacharya, Bette Korber, and Will Fischer. Web-based design and evaluation of T-cell vaccine candidates. Bioinformatics, 24(14):1639–1640, July 2008.

[40] Kazutaka Katoh, John Rozewicki, and Kazunori D. Yamada. MAFFT online service: Multiple sequence alignment, interactive sequence choice and visualization. Briefings in Bioinformatics, June 2017.

[41] Régine Audran, Michel Cachat, Floriana Lurati, Soe Soe, Odile Leroy, Giampietro Corradin, Pierre Druilhe, and François Spertini. Phase I Malaria Vaccine Trial with a Long Synthetic Peptide Derived from the Merozoite Surface Protein 3 Antigen. Infection and Immunity, 73(12):8017–8026, December 2005.

[42] Sheetij Dutta, Lisa S. Dlugosz, Damien R. Drew, Xiaopeng Ge, Xiopeng Ge, Diouf Ababacar, Yazmin I. Rovira, J. Kathleen Moch, Meng Shi, Carole A. Long, Michael Foley, James G. Beeson, Robin F. Anders, Kazutoyo Miura, J. David Haynes, and Adrian H. Batchelor. Overcoming antigenic diversity by enhancing the immunogenicity of conserved epitopes on the malaria vaccine candidate apical membrane antigen-1. PLoS pathogens, 9(12):e1003840, 2013.

[43] Wenling Wang, Renqing Li, Yao Deng, Ning Lu, Hong Chen, Xin Meng, Wen Wang, Xiuping Wang, Kexia Yan, Xiangrong Qi, Xiangmin Zhang, Wei Xin, Zhenhua Lu, Xueren Li, Tao Bian, Yingying Gao, Wenjie Tan, and Li Ruan. Protective Efficacy of the Conserved NP, PB1, and M1 Proteins as Immunogens in DNA- and Vaccinia Virus-Based Universal Influenza A Virus Vaccines in Mice. Clinical and vaccine immunology: CVI, 22(6):618–630, June 2015.

[44] Suzanne L. Epstein, Terrence M. Tumpey, Julia A. Misplon, Chia-Yun Lo, Lynn A. Cooper, Kanta Subbarao, Mary Renshaw, Suryaprakash Sambhara, and Jacqueline M. Katz. DNA Vaccine Expressing Conserved Influenza Virus Proteins Protective Against H5N1 Challenge Infection in Mice. Emerging Infectious Diseases, 8(8):796–801, August 2002.

[45] Annette von Delft, Timothy A. Donnison, José Lourenço, Claire Hutchings, Caitlin E. Mullarkey, Anthony Brown, Oliver G. Pybus, Paul Klenerman, Senthil Chinnakannan, and Eleanor Barnes. The generation of a simian adenoviral vectored HCV vaccine encoding genetically conserved gene segments to target multiple HCV genotypes. Vaccine, 36(2):313–321, January 2018.

[46] A. H. Land and A. G. Doig. An automatic method of solving discrete programming problems. Econometrica, 28(3):497, July 1960.

[47] Charles Audet and Dominique Orban. Finding Optimal Algorithmic Parameters Using Derivative-Free Optimization. SIAM Journal on Optimization, 17(3):642–664, jan 2006.

[48] Mustafa Baz, Brady Hunsaker, and Oleg Prokopyev. How much do we “pay” for using default parameters? Comput Optim Appl, 48(1):91–108, jan 2011.

[49] Frank Hutter, Holger H. Hoos, and Kevin Leyton-Brown. Automated Configuration of Mixed Integer Programming Solvers. In Andrea Lodi, Michela Milano, and Paolo Toth, editors, Integration of AI and OR Techniques in Constraint Programming for Combinatorial Optimization Problems, Lecture Notes in Computer Science, pages 186–202. Springer, 2010.

