## Supplement One for "Graph-Theoretical Formulation of the Generalized Epitope-based Vaccine Design Problem"

### A Obtaining Pareto-efficient solutions in a multi-objective optimization problem

We wish to explore the Pareto frontier of this bi-objective optimization problem (for simplicity, we assume there is only a single subset $P$):

$$\begin{aligned} \text{Maximize} \quad & z = \left\{ I(P) - \sum_{e_{ij} \in H(P)} w(e_{ij}) \right\} \\ \text{Subject to} \quad & |P| \leq k \end{aligned} \tag{1}$$

One possible algorithm to do so is the augmented $\epsilon$-constraint method [1], an iterative solving approach in which one of the two objectives is moved into the constraints and bounded to take values smaller than $e_i$:

$$\begin{aligned} \text{Maximize} \quad & I(P) - \epsilon s \\ \text{Subject to} \quad & \sum_{e_{ij} \in H(P)} w(e_{ij}) - s = e_i \\ & |P| \leq k, s \in \mathbb{R}^+ \end{aligned} \tag{2}$$

with $\epsilon$ small. The surplus variable $s$ is used in the objective function to discard weakly efficient solutions, when multiple optimal solutions with the same immunogenicity exist. The bounds $e_1, \ldots, e_N$ uniformly span the range between

$$\min_{|P| \leq k} \sum_{e_{ij} \in H(P)} w(e_{ij}) \tag{3}$$

and

$$\min_{|P| \leq k} \sum_{e_{ij} \in H(P)} w(e_{ij}) \text{ s.t. } I(P) = \max_{|P| \leq k} I(P) \tag{4}$$

i.e. the minimum edge cost, and the minimum edge cost with the maximum unconstrained immunogenicity. This procedure requires to solve $N + 3$ instances of the problem and results in $N$ points on the Pareto frontier balancing the immunogenicity and the cumulative cleavage likelihood.

### B Measuring epitope conservation in terms of entropy

We quantified the variation in each position of the pathogen by computing the entropy on aligned sequences, only in the position where the consensus was not a gap. [2] showed that the entropy at a given position is an acceptable metric to quantify residue conservation, and, indirectly, its importance for the structure and function of the protein. The entropy of the $i$-th residue is computed as:

$$H_i = -\sum_a \frac{n_{ai}}{N} \log\left(\frac{n_{ai}}{N}\right) \tag{5}$$

with $n_{ai}$ counting how many aligned sequences contain the amino acid $a$ in position $i$, and $N$ the number of sequences. Since we are interested in the conservation of 9-mers, we smooth the entropy using a triangular filter of width 9:

$$\tilde{H}_i = \frac{1}{25} \sum_{j=-4}^{4} (5 - |j|) H_{i+j} \tag{6}$$

### References


[1] George Mavrotas. Effective implementation of the $\epsilon$-constraint method in Multi-Objective Mathematical Programming problems. *Applied Mathematics and Computation*, 213(2):455–465, July 2009.

[2] William S.J. Valdar. Scoring residue conservation. *Proteins: Structure, Function, and Genetics*, 48(2):227–241, August 2002.
