## Supplement Two for "Graph-Theoretical Formulation of the Generalized Epitope-based Vaccine Design Problem"

| | | |
|---|---|---|
| Maximize | | |
| (OBJ) | $\sum_{t\in t}\sum_{v\in v}\sum_{a\in a} y_{vt}p_a i_{va} + \sum_{o\in o}\theta_o$ | |
| Subject to | | |
| (C1) | $\sum_{v\in v} x_{wvt} = \sum_{v\in v} x_{vwt} = y_{wt}$ | $\forall w \in v, t \in t$ |
| (C2) | $\sum_{t\in t} y_{vt} \leq 1$ | $\forall v \in v$ |
| (C3) | $\sum_{t\in t}\sum_{v\in v} x_{svt} = \sum_{t\in t}\sum_{v\in v} x_{vst} = \lvert t\rvert$ | |
| (C4) | $\sum_{(v,w)\in e} x_{vwt} w(e_{vw}) \leq h$ | $\forall t \in t$ |
| (C5) | $\sum_{v\in v} y_{vt} \leq k$ | $\forall t \in t$ |
| (C6a) | $u_{vt} - u_{wt} + 1 \leq (\lvert v\rvert - 1)(1 - x_{vwt})$ | $\forall e_{vw} \in e, t \in t$ |
| (C6b) | $1 \leq u_{vt} \leq \lvert v\rvert - 1$ | $\forall v \in v, t \in t$ |
| (C7) | $\sum_{t\in T}\sum_{v\in V} y_{vt}\tau_{vo} \geq \theta_o$ | $\forall o \in O$ |
| (C8) | $x_{vwt}, y_{vt}, \theta_o \in \{0, 1\}$ | $\forall v, w, t, o$ |

Table S2: Alternative ILP formulation where pathogen coverage is maximized together with immunogenicity.

| | | |
|---|---|---|
| Maximize | | |
| (OBJ) | $\sum_{t\in t}\sum_{v\in v}\sum_{a\in a} y_{vt} p_a i_{va} + \sum_{t\in T}\sum_{v\in V}\sum_{o\in O} y_{vt}\tau_{vo}$ | |
| Subject to | | |
| (C1) | $\sum_{v\in v} x_{wvt} = \sum_{v\in v} x_{vwt} = y_{wt}$ | $\forall w \in v, t \in t$ |
| (C2) | $\sum_{t\in t} y_{vt} \leq 1$ | $\forall v \in v$ |
| (C3) | $\sum_{t\in t}\sum_{v\in v} x_{svt} = \sum_{t\in t}\sum_{v\in v} x_{vst} = \|t\|$ | |
| (C4) | $\sum_{(v,w)\in e} x_{vwt} w(e_{vw}) \leq h$ | $\forall t \in t$ |
| (C5) | $\sum_{v\in v} y_{vt} \leq k$ | $\forall t \in t$ |
| (C6a) | $u_{vt} - u_{wt} + 1 \leq (\|v\| - 1)(1 - x_{vwt})$ | $\forall e_{vw} \in e, t \in t$ |
| (C6b) | $1 \leq u_{vt} \leq \|v\| - 1$ | $\forall v \in v, t \in t$ |
| (C7) | $x_{vwt}, y_{vt} \in \{0, 1\}$ | $\forall v, w, t$ |

Table S3: Alternative ILP formulation where epitope conservation is maximized together with immunogenicity.
