## Supplement Three for "Graph-Theoretical Formulation of the Generalized Epitope-based Vaccine Design Problem"

| Allele | Freq. | Allele | Freq. | Allele | Freq. |
|---|---|---|---|---|---|
| A*01:01 | 4.50 | A*02:01 | 10.69 | A*02:05 | 0.88 |
| A*03:01 | 3.69 | A*11:01 | 7.52 | A*24:02 | 12.91 |
| A*31:01 | 2.43 | A*68:01 | 1.77 | B*07:02 | 3.61 |
| B*08:01 | 2.95 | B*15:01 | 2.06 | B*27:02 | 0.15 |
| B*27:05 | 1.11 | B*35:01 | 3.24 | B*37:01 | 0.44 |
| B*38:01 | 0.66 | B*39:01 | 1.77 | B*40:01 | 5.31 |
| B*40:06 | 0.52 | B*44:03 | 2.21 | B*51:01 | 3.24 |
| B*51:02 | 0.22 | B*52:01 | 0.88 | B*58:01 | 2.65 |
| C*04:01 | 8.26 | C*06:02 | 5.09 | C*07:02 | 9.66 |

Table S1: HLA alleles and their percent frequency in the world population used in this study.
